# Synthesis of Extended, Self-Assembled Biological Membranes containing Membrane Proteins from Gas Phase

**DOI:** 10.1101/661215

**Authors:** Matthias Wilm

## Abstract

Membrane proteins carry out a wide variety of biological functions. The reproduction of their specific properties in a technically controlled environment is of significant interest. Here, a method is presented that allows the self-assembly of a macroscopically large, freely transportable membrane with Outer membrane porin G from Escherichia Coli. The technique does not use protein specific characteristics and therefore could represent a path to the generation of extended layers of membranes with integrated, arbitrary membrane proteins. The composition of the membrane, its lipid and protein content, is experimentally controlled. Such in-vitro systems are relevant for the study of membrane-protein function and structure and the self-assembly of membrane-based protein complexes. They might become important for the incorporation of lipid-membranes into technical devices.

## I. Introduction

This article relates to two separate scientific domains, that of nanoelectrospray based mass spectrometry and membrane research. The author has a background in electrospray based mass spectrometry. Any imperfections in terminology and lack of rigor in the field of membrane biology may be excused.

The purpose of the manuscript is to show experimental evidence that nanoelectrospray can be used to synthesize an environment in which large membranes with integrated membrane proteins can self-assemble. The overall experimental set-up follows a common technique used in solid-state-physics and applies it to a biological context: the synthesis of novel materials by chemical vapor deposition (for instance in “Thin Solid Films”, Elsevier).

The article is divided into two separate parts. The first showing the experiments that led to the conclusion that a nanoelectrospray, when operated correctly, is a vapor deposition device for large molecules. And the second demonstrating how it can be used to create an environment in which membranes self-assemble.

## II. Nanoelectrospay as a Molecular Beam Device

### A. Introduction

With the discovery of soft-ionization methods Matrix Assisted Laser Desorption/Ionization (MALDI) and electrospray by Hillenkamp and Fenn respectively in the late 80ties it became evident that large molecules can be stably transferred into gas phase (1–3). Less known is that the first nanoelectrospray device, a perfect electrospray ion source inspired by the theoretical description of the electrospray process, was built at the same time in the physics department of just the university where MALDI was being discovered (4). It could be shown that this device was able to generate molecular mono-layers of a non-volatile substance when spraying a solution of it onto a surface. The so far unpublished results of the mono-layer experiments from 1988/89 are presented here because they form the scientific base of the membrane synthesis method.

### B. Nano Electrospray Based Surface Preparation

In 1988, before the discovery that large molecules can be stably transferred into gas phase, the author built an electrospray device for the homogenous distribution of a not volatile substance over a metallic surface. This was done as an improved sample preparation method for an investigation with a static static organic Time-of-Flight-Secondary Ion Mass Spectrometry (TOF-SIMS) instrument (5). Static organic SIMS is capable of analyzing molecules on surfaces with submonolayer sensitivity. Ar^+^ ions, hitting a metallic target, transfer their kinetic energy into the surface (5). Organic molecules lying on the surface detach by partially absorbing this kinetic energy. They acquire a charge by ion-capture or by their intrinsic ionic state and accelerate away from the target pulled by strong electric fields. A time-of-flight mass spectrometer measures their mass/charge ratio. The desorption mechanism explains the sub-monolayer sensitivity of static SIMS. If the surface is covered by more than one layer, the uppermost molecules do not absorb sufficient energy from the metal target. They cannot desorb and the static SIMS signal decreases. At the time, before the discovery of the Matrix Assisted Laser Desorption/Ionization (MALDI) and electrospray ionization processes, static SIMS was one of the most sensitive techniques for the analysis of organic molecules of up to 10 kDa in size.

Conventional electrosprays generate thin films for many different purposes (6). In general particles in the μm size range are distributed homogeneously over the prepared surface (see figure 1). The first electrospray apparatus built reproduced this behavior. It operated at conventional flow rates of 10 - 20 μl/ min. Figure 2 panel A shows the scanning electron microscopic picture of a surface prepared with an acetone solution of rhodamine B using this device. After the acetone evaporated in flight rhodamine particles with sizes between 0.2 μm and 1.5 μm cover the aluminum surface. The rhodamine on the surface was visible as a white homogeneous layer.

**Fig. 1.**
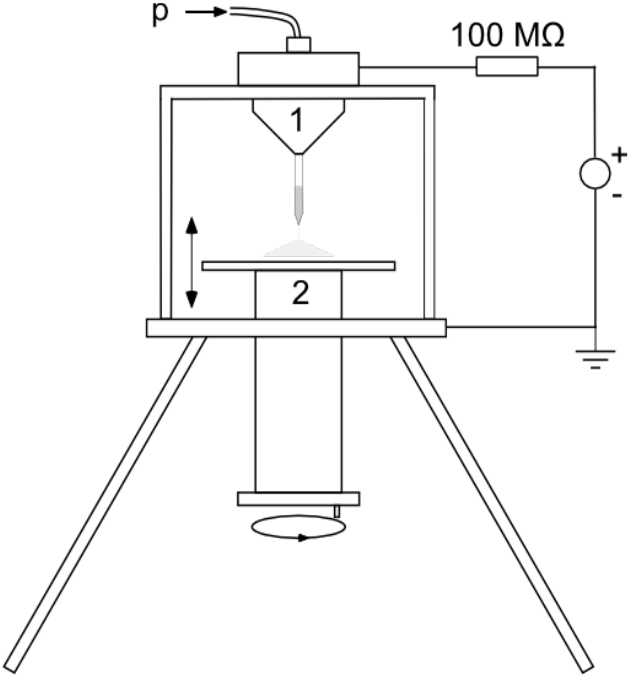
Electrospray Apparatus for Surface Preparation. The electrospray apparatus for the preparation of membranes (2nd generation instrument): a pressurized container holds a gold-coated pulled glass capillary filled with 1 or 2 μl solution (1). An optional small pressure supports the continuous flow of the sample through the glass capillary during operation. The distance between the grounded target (2) and the needle is adjustable. It was usually 3 cm. The 1st generation instrument had a 250 μm steel nozzle and, after that, a manually pulled glass capillary as an emitter. Its container was not pressurized.

**Fig. 2.**
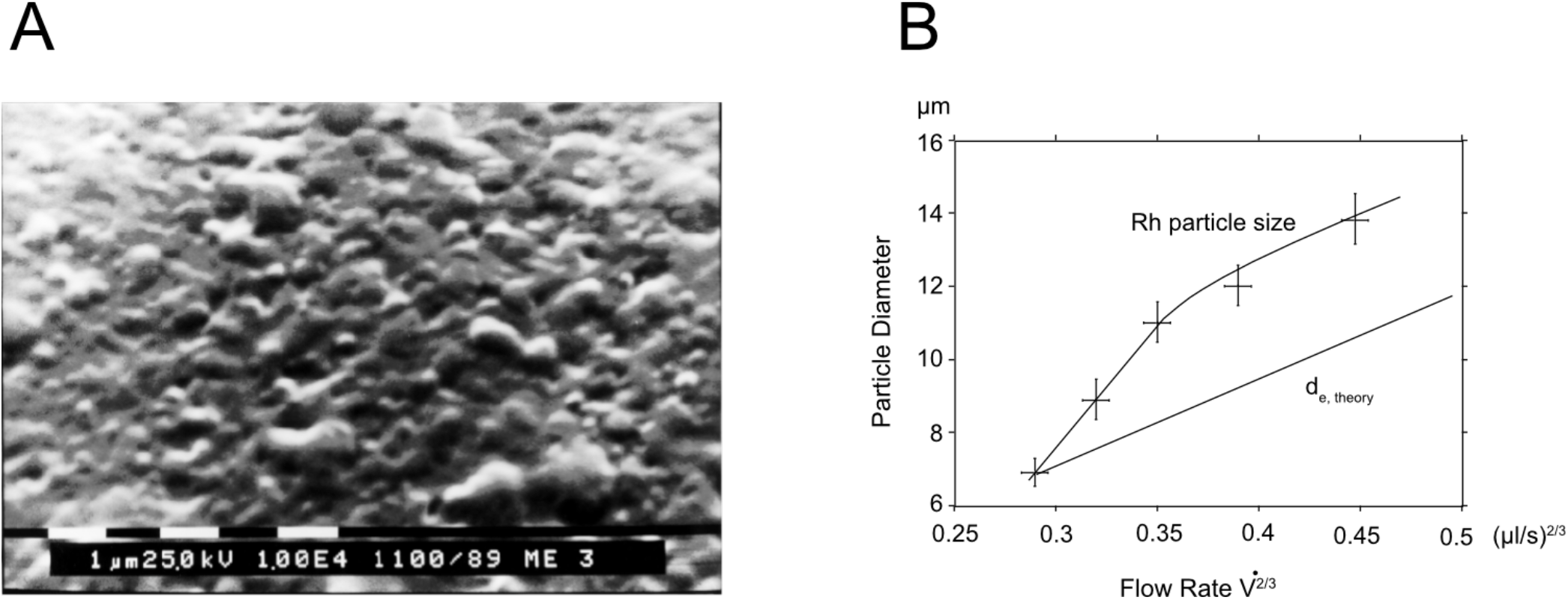
Conventional Electrospray Prepared Surface. Panel A shows the raster-electron microscopic picture of a surface prepared with an electrospray from a 250 μm nozzle. The surface layer originated from a 30 μM acetone solution sprayed onto a metal target with a flow rate of 8 - 15 μl/min. The image shows a 10 000x magnification of the gold coated rhodamine surface. The surface is densely covered with 0.2 μm - 1.5 μm sized particles. Panel B shows that the maximal observable particle size on the target depends on the flow rate in agreement with formula 1. When spraying a saturated rhodamine/acetone solution, the particle diameter on the target corresponds roughly to the size of the initially produced droplets.

Such a thick preparation is insufficient for a static SIMS investigation which analyses sub-monolayer structures. Theoretical and experimental investigations into the electrospray process itself showed a way how to improve the homogeneity of the preparation (7). A stable liquid Taylor cone ejects droplets exclusively from its tip. The radius for the emission of droplets depends on the flow rate. The central equation describing the radius of the emission region from the tip of a Taylor cone is (7):

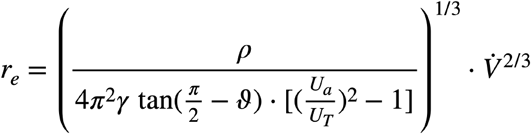

*r_e_*: emission radius of droplets at the tip of the Taylor Cone,
*ρ*: density of the solution, *γ*: surface tension,
*ϑ*: opening angle of the Taylor Cone (49.3° in the static case),
*U_a_*: applied and *U_T_* Taylor Cone threshold voltage,
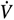: flow rate.

**Equation 1**

The central equation linking the radius re of the droplet emission region at the tip of a Taylor cone with the flow rate V′.

Reducing the flow rate reduces the size of the initially produced droplets (figure 2, panel B).

The equation suggests that when spraying an aqueous solution of 1 pmol/μl at a flow rate of 20 nanoliters/min, droplets with the size of the emission radius would contain, on average, only one analyte molecule.

Acetone has a high vapor pressure. Its droplets start to evaporate in flight and break up into smaller second generation droplets before they dry down and the non volatile residuals reach the surface (8). The droplet emission radius is only a description for the largest droplet generated by the spray, not the most common one. By replacing the metallic nozzle with a gold-coated, pulled glass capillary, the acetone flow rate was about 200 nanoliters/min. This was the lowest flow rate I could achieve at the time with the first generation instrument. With this emitter, the metallic aluminum surface did not visibly change when the rhodamine solution was sprayed - even under prolonged preparation of 1 h or more. A raster-electron microscope and a static SIMS mass spectrometer analyzed the obtained targets (figure 3, panel A, B). The raster electron microscopic picture showed no discernible particles even though rhodamine covered the surface of the target.

**Fig. 3.**
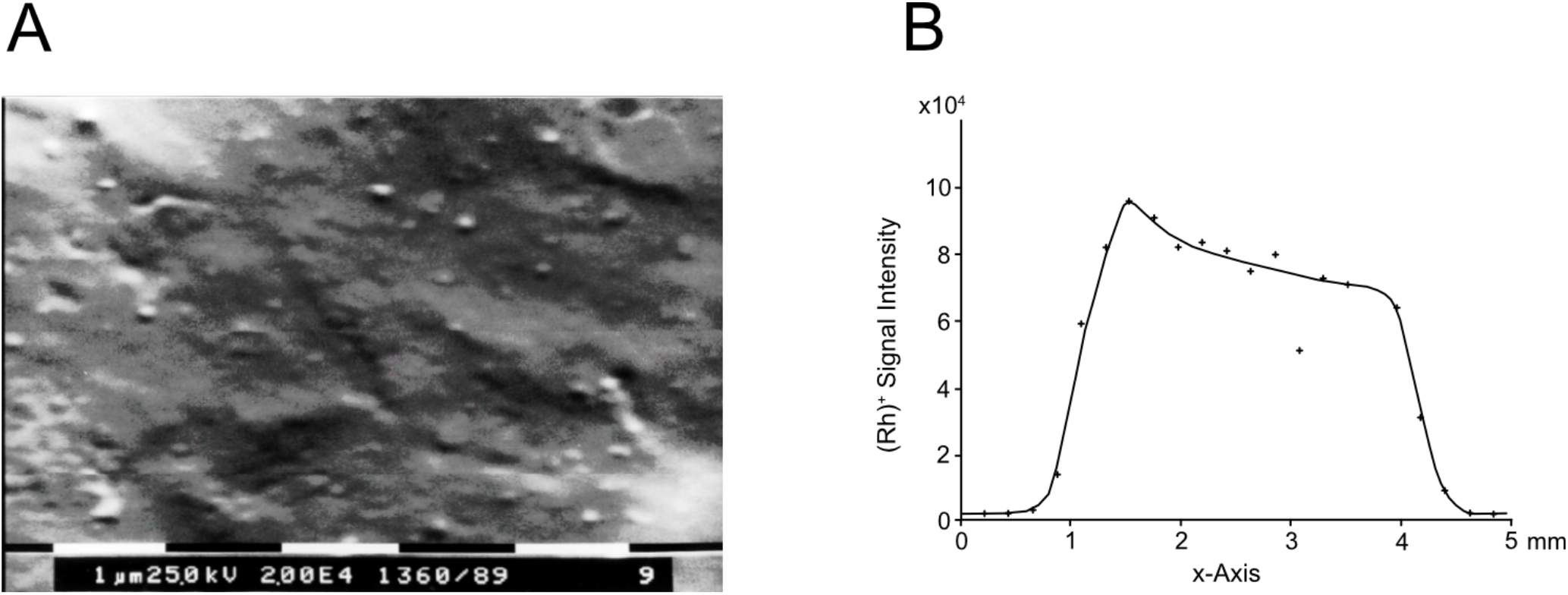
Nanoelectrospray Prepared Surface. Panel A shows a rhodamine prepared layer using a self-sustained flow rate of approximately 200 nanoliters/min generated with a hand-pulled gold-coated glass capillary. The photo shows the 20 000x magnified surface. While rhodamine covered the entire surface (see panel B), no specific particle sizes could be determined. Panel B shows the static SIMS signal of the same target as in panel A, only before it was coated with gold. The spatially resolved static SIMS investigation showed that the rhodamine covered a 4 mm wide area evenly. At the time, this target represented the most homogeneously prepared surface ever investigated on this machine (9) (TOF-SIMS II 9, https://www.iontof.com/history-ion-tof-sims.html).

The following experiments were intended to find out whether the nanoelectrospray was able to prepare rhodamine monolayers. Such a preparation would mean that the ultimately generated droplets were so small that they contained on the average only one rhodamine molecule. The experimental results achieved at this stage were not completely conclusive but they provided enough evidence to continue on this path of experimentation.

Figure 4 panel A shows the slow build up of the static SIMS signal with preparation time and a fall off when the rhodamine coverage becomes too thick. This behavior is similar to a monolayer coverage on clean surfaces in vacuum by molecular beam deposition (5). The question was whether the surface is covered by an increasing number of particles, as in panel A of figure 2, or if the coverage is monolayer-like. To answer this question, a series of experiments determined the time required to reach the maximum SIMS response signal t_SIMSmax_ as a function of the rhodamine concentration. If small rhodamine particles cover the target t_SIMSmax_ depends on their cross-section, hence varies with (c_Rh_)^-2/3^. However, if the coverage is monolayer-like, the static SIMS signal responds to the total amount of rhodamine sprayed onto the surface and will depend on (c_Rh_)^-1^. Figure 4 panel B shows how the preparation time to reach the maximum SIMS signal t is indirectly proportional to the rhodamine concentration. The linearity indicates that the SIMS signal characteristic changes proportionally to the total amount of rhodamine sprayed onto the target. This result is an indication that the surfaces generated by the nanoelectrospray preparation behave like molecular monolayers under a static SIMS investigation.

**Fig. 4.**
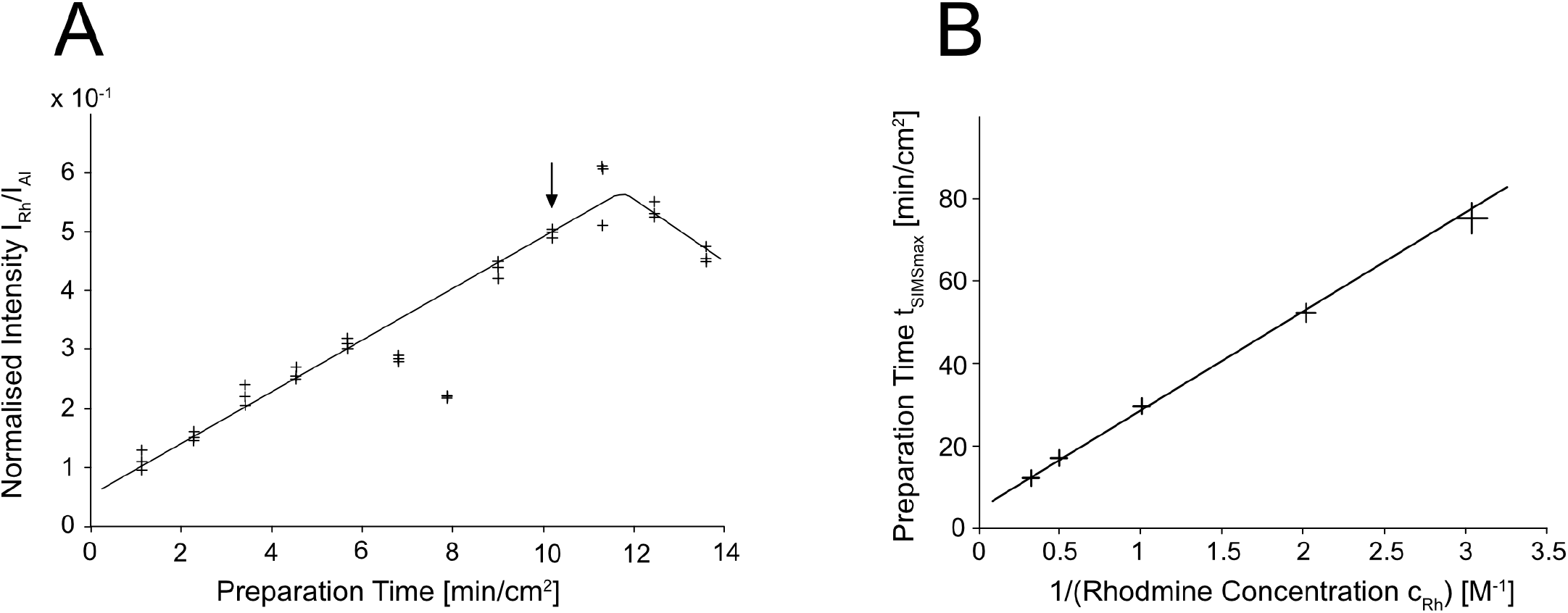
Built up of the Rhodamine Coverage with Nanoelectrospray Prepration Time. Static SIMS response characteristic as a function of the amount of rhodamine sprayed onto the surface: panel A shows the increasing Rh^+^ signal with preparation time. The sprayed liquid was a 30 μM rhodamine-acetone solution. For each target, three different measurements at different target locations contribute to the analysis. The SIMS signal intensity rises with increasing surface coverage passes through a maximum, and drops off. This characteristic is similar to surfaces which are successively covered by molecular beams in vacuo. During the preparation of targets 6 and 7, the electrospray became temporarily unstable. A more thorough analysis with target 9 using spatially resolved SIMS and electron microscopy revealed more details of the surface topology (figure 3, Panel A, B). Panel B shows the relationship between the time taken to reach the maximum SIMS response signal t_SIMSmax_ and the rhodamine concentration in the sprayed solution. This time is indirectly proportional to the rhodamine concentration.

#### Molecular Beam Hypothesis

The surface preparation experiments concluded in that when an acetone solution is electro-sprayed from a stable Taylor cone using a metal coated pulled glass capillary with a flow rate in the order of 200 nanoliters/min the compartmentalization of individual analyte molecules into separate droplets is complete or close to complete (8). Two hypothesis arose from these experiments. First: if such a device is directed towards a vacuum system, the volatile solvent would evaporate and release the non volatile molecule into the vacuum. Like this a nanoelectrospray is an ion source for large molecules of virtually unlimited mass - that, what electrospray ion sources are today. And second: it should be possible to use a nanoelectrospray to synthesize molecularly designed layers of large organic compounds (4).

While these experiments were in its final stage, Prof. Fenn et al. published the first mass spectra of protein ions generated with a conventional electrospray ion source (1; 2).

In 1992 the first project was realized with the implementation of the nanoelectrospray ion source at the European Molecular Biology Laboratory (EMBL)(10). The direct interfacing of the nanoelectrospray ion source with the vacuum system of a mass spectrometer demonstrated that the compartmentalization of analyte molecules is indeed complete. The nanoelectrospray ion source was about 100 times more sensitive than any other electrospray ion source at the time. The device provided the technical base for the development of low-level protein identification by mass spectrometric means. Over the years it was used to analyze thousands of peptide and protein samples.

**Fig. 5.**
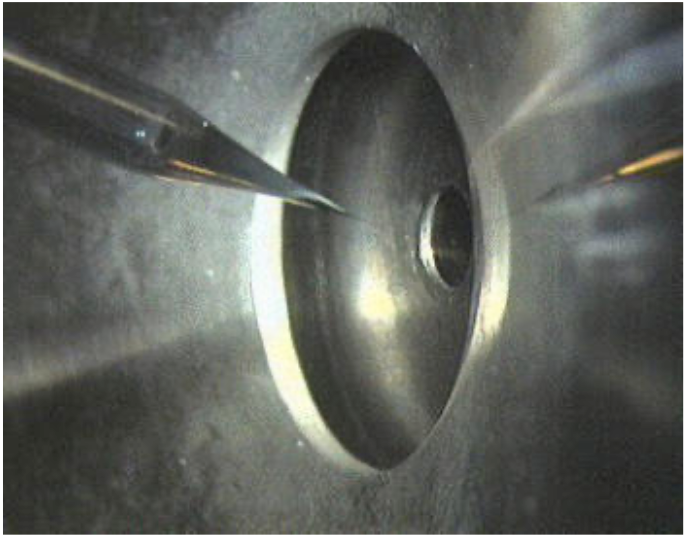
Nanoelectrospray Ion Source on a Mass Spectrometer. The nanoelectrospray ion source mounted on a triple quadrupole mass spectrometer (API III, Sciex). The image shows the tip of the emitter which is mounted in the same pressurized holder as in figure 1. The pulled glass needle is placed centrally directly in front of the orifice to the vacuum system of the mass spectrometer. A flight path of about 1 cm under atmospheric conditions is sufficient to allow complete desolvation of large ions from aqueous solutions. No protein clusters are present in the mass spectra.

## III. Self Assembly of Large Membranes

Can a nanoelectrospray ion source be used to synthesize membranes with incorporated membrane proteins from gas phase ?

Because purified membrane proteins are only stable when associated with detergent molecules, it is not possible to synthesize such membranes directly (11; 12). The question was whether nanoelectrosprayed layers could constitute an environment in which they self-assemble. For this, the ability of the nanoelectrospray to generate mono-molecular films was the experimental foundation. The issue to address was: what are the requirements for self-assembly of protein filled lipid bilayers when synthesized in a layered fashion? The course of the experiments suggests the answer: hydrostatic stabilization of the membrane proteins on both sides of the lipid bilayer. Even though a lipid bilayer can form on the surface of a liquid, incorporation of the proteins will only occur when a hydrostatic layer thick enough to enclose the proteins surrounds the lipid layer. This experimental finding corresponds to the observation that membrane proteins in a supported bilayer can denature when the distance between the membrane and the underlying support is too small (13).

The experimental results presented here are not sufficient to unequivocally prove this claim but similar to the experiments done in 1988/89 they provide very strong evidence that it was indeed possible to generate a very large membrane with incorporated membrane proteins. Enough to justify additional experimental efforts.

The first experimental result was the observation that lipid mono- and bilayers can assemble on a liquid surface when the lipids are sprayed onto them. The substrate selected for the layer generation was the liquid meniscus of a small round container with a diameter of 3 mm corresponding to the size of a standard electron microscopic grid. This is a surface area of 7.07 mm^2^. When distributing 33 pmols phosphatidylcholine evenly over the surface of the meniscus the lipid density is 36 Å^2^ per molecule. This value lies between the surface occupancy of 60 Å^2^ - 70 Å^2^ as measured for phosphatidylcholine molecules in vesicles (14), and approximately 20 Å^2^ for a saturated fatty acid molecule in a regular Langmuir-Blodgett film (15). Considering the linear structure of saturated fatty acids and their dense packing in a regular array, 20 Å^2^ is certainly too small for the surface occupancy of a phosphatidylcholine molecule in a closed, flat phosphatidylcholine film. When the layer consisted of 66 pmols phosphatidylcholine, the surface viscosity visibly changed, and, after overnight incubation, folds in the surface appeared (see Figure I, supplemental material). The folding did not occur when the layer contained only 33 pmols. This indicates that a membrane had formed.

The behavior of outer membrane protein G (ompG) from Escherichia Coli on the surfaces generated by spraying 66 pmols and 33 pmols underlines their different nature. ompG is a relatively resilient 35 kDa monomeric channel protein. It is a membrane spanning protein that denatures in an aqueous buffer if no detergent is present to cover its hydrophobic regions. Its crystal structure shows that it has an outer pore diameter of approximately 2.5 nm in the open conformation with a 2 nm large central channel at its periplasmatic side (11; 12). The observation was that ompG sprayed onto a surface prepared with 66 pmols phosphatidylcholine denatures upon removal of the detergent. When sprayed onto a surface with only 33 pmols phosphatidylcholine it remains intact.

Figure 6 shows transmission-electron microscopic images of the first successfully assembled membrane layer containing ompG. These images come from an early stage of successful experiments. The preparation protocol was subsequently optimized to generate more extended protein-lipid membranes in a more reproducible way. However, they demonstrate that no support other than the liquid surface is required to allow membrane assembly.

**Fig. 6.**
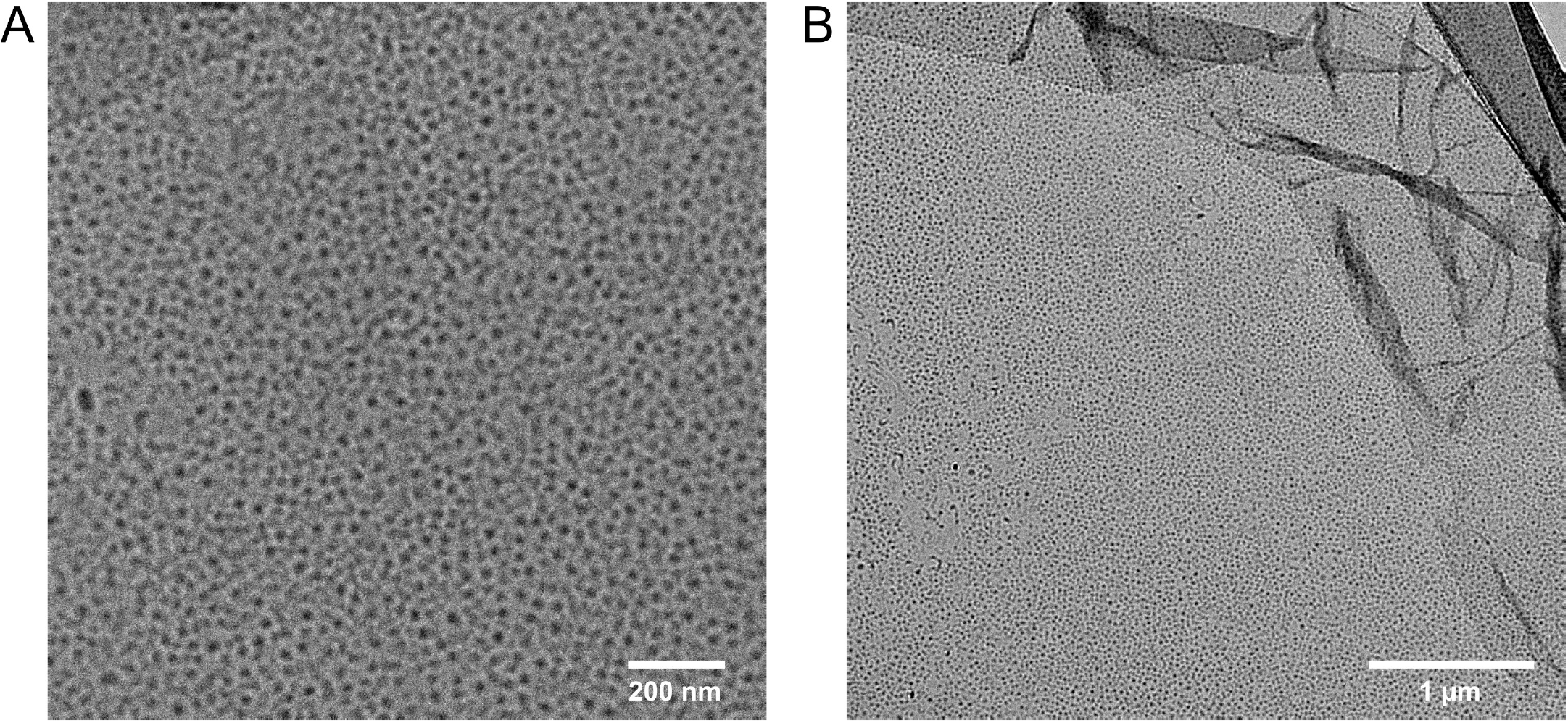
ompG Containing Membrane on Buffer Solution. Transmission electron microscopic picture of the first successfully self-assembled ompG containing membrane prepared with nanoelectrospray onto a buffer solution: a negative stain with a 1% uranyl-acetate solution generates the contrast. The folded structure in panel B is the ruptured and crumbled carbon layer of the EM grid at this location.

The first step for this preparation was to spray 1 μl 22 pmol/ μl phosphatidylcholine in ethanol onto the liquid container holding a 10 mM TRIS solution, pH 7.5 and some SM2-Biobeads. Then followed the distribution of 1 μl 16 pmol/μl ompG in 1 w% octylglucoside 10 mM TRIS solution and again 1 μl 22 pmol/μl phosphatidylcholine. The fourth layer consisted of glycerol by spraying approximately 3 μl 1:2:2 solution of glycerol, water, and ethanol. The container was incubated at 37 °C in a closed petri dish in a saturated water vapor atmosphere for seven days to allow the extraction of the octylglucoside detergent by the SM2-Biobeads. Then, the liquid surface was transferred to a carbon-coated copper grid, washed several times with doubly distilled water, negatively stained with 1% uranyl acetate and imaged in an transmission electron microscope.

One particular feature of the image came as a surprise - the electron dense center of the pores. A possible explanation for this is that the washing steps failed to remove glycerol from the interior of the pores. Without additional measures, liquid water cannot readily permeate pores with a diameter in the low nm range. This aspect is central to Gore-Tex membranes which are permeable to gaseous but not to liquid water. Uranyl-acetate staining tends to enlarge features in EM pictures. To acquire images that represent the layer more accurately, we chose platinum shadowing for subsequent ompG analyses.

The preparation protocol was further optimized to generate large membrane layers in a more reproducible and robust way (figure 7). The first step consists of spraying 66 pmols of bipolar lipid onto a buffer containing SM-2 biobeads. The container was incubated overnight at room temperature. This generates a bi-lipid layer that retains subsequent molecules in the surface. A thin layer of glycerol laid the foundation for the membrane assembly in a hydrophilic environment by spraying a 1:2 glycerol/ethanol solution. The equivalent of a lipid mono-layer was added (33 pmols) followed by the detergent-solution of ompG and sufficient lipid to form a closed bilayer (20 - 30 pmols). A final glycerol layer completes the hydrophilic environment.

**Fig. 7.**
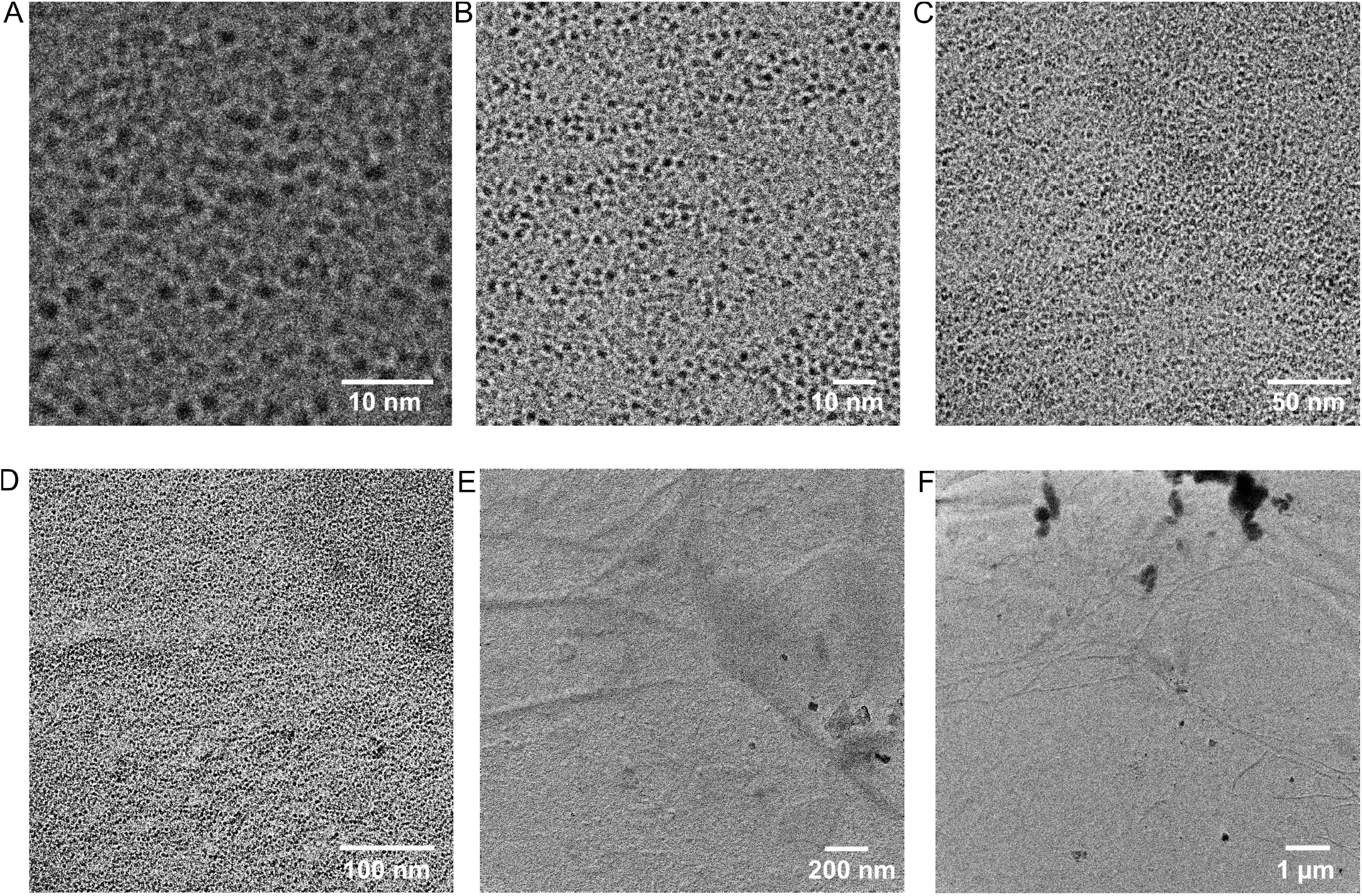
Large ompG Containing Membrane prepared by Nanoelectrospray in a Layered Fashion. The images show a transmission-electron microscopic investigation of an ompG containing membrane prepared with nanoelectrospray and shadowed with a 1.5 nm thick platinum layer at an angle of 15°. The images display central zoom outs from the highest resolved image in panel A to capture the self-similarity of the homogeneous layer. A monolayer of the ompG filled membrane covers the entire region. Such a free floating membrane has never been seen before.

The SM-2 beads in the buffer extracted the detergent from the surface in a six days lasting incubation period. After transfer to a EM grid and washing with distilled water the grid was exposed to a platinum beam in vacuo. Figure 7 shows a series of images from one region of a lipid bilayer formed from this assembly. A single, extended, intact ompG containing layer has formed. No denatured protein was visible. ompG denatures in an aqueous environment when not stabilized by detergent or lipids. The presence of the structures depends on the addition of ompG to the surface, and they are virtually identical in size and shape to a projection of the space-filling model of ompG calculated from crystallization data (12) (see Movie I, supplemental material).

## IV. Protein Complex Formation in Nanoelectrospray Generated Layers

The assembly of a large protein containing membrane proceeds in a thin glycerol layer which most likely too thin to promote the formation of membrane stacks or tubes. The question is whether this highly artificial environment allows the assembly of membrane based protein complexes. Even though this has to be decided on a case by case basis we tested it on the proteins listeriolysin O and pneumolysin. Listeriolysin O and pneumolysin are both members of a group of pore-forming toxins. They are soluble 53 kDa respective 60 kDa proteins. Up to 50 monomers build a non-covalent, ring-shaped complex on membranes. The rings insert themselves and create large pores of about 30 nm in diameter (16; 17). Listeriolysin O assembles best under a slightly acidic pH of 5.5. Both proteins require phospholipids in the membrane for docking and complex formation.

### A. Listeriolysin O Assembly on a Lipid Bilayer

The buffer chosen for the preparation of listeriolysin O layers allows the assembly of its complexes on membranes (18). However, in these experiments here, the local environment for the protein is a thin glycerol film and not the buffer itself. In thin layers, ion concentrations and pH can be very different from bulk solution. In an experiment to test the complex formation, the nanoelectrospray generated first a lipid bilayer film, then a listeriolysin O containing film followed by a glycerol layer to finalize the hydrophilic environment. Figure 8 shows the result. The overall appearance corresponds to published listeriolysin O complexes on membranes (18). The complex assembly is highly efficient. Again, the electron-dense centers of many of the completed rings are probably attributable to remnants of glycerol and salt residues from the buffer solution that the washing procedure did not successfully remove. Mono-molecular ompG pores in a nanoelectrospray prepared layer have a similar appearance (see figures 6, 7).

**Fig. 8.**
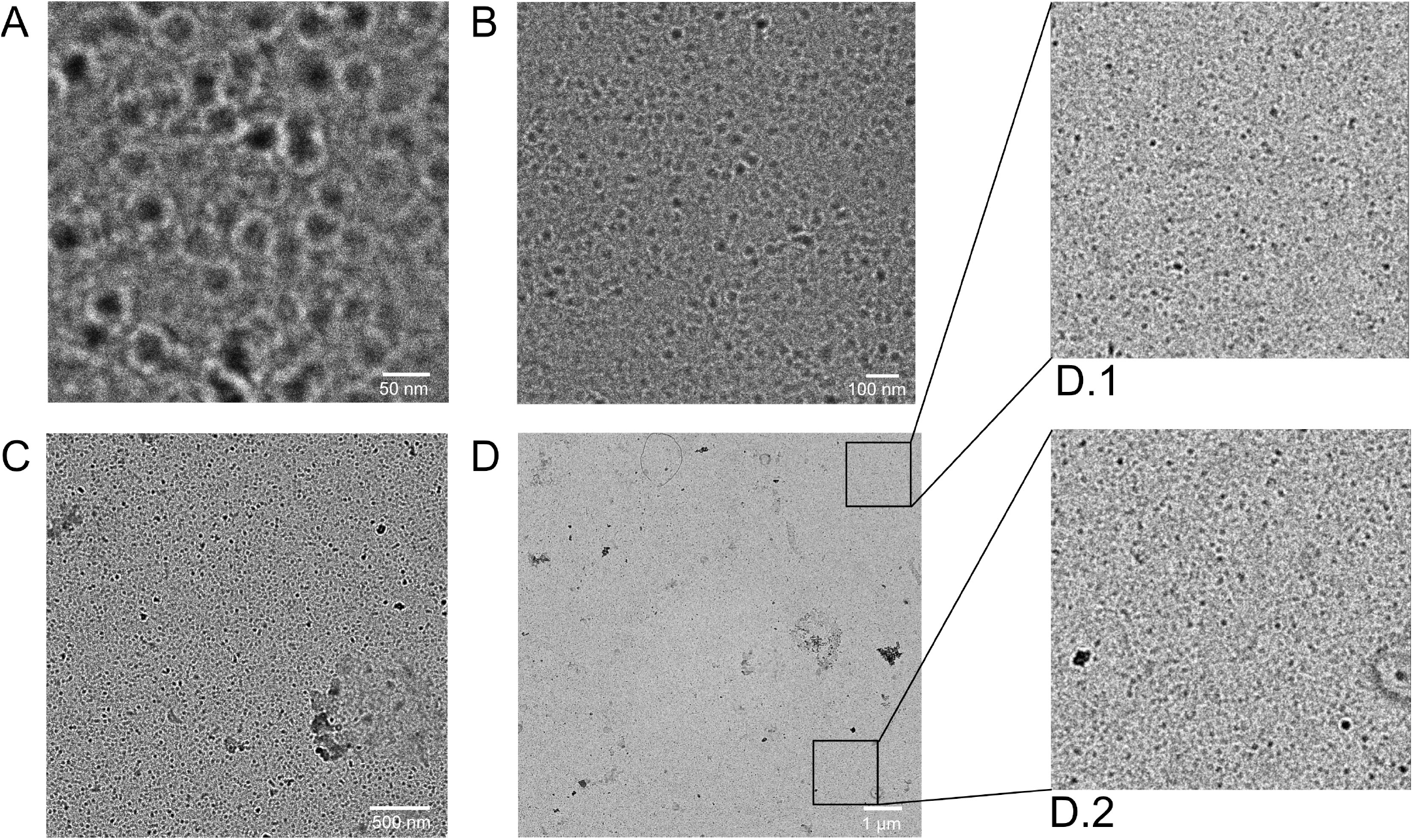
Listeriolysin O Assembly on a Lipid Bilayer. Transmission electron microscopic image of listeriolysin O sprayed onto a lipid bilayer and covered with glycerol: A negative stain with 1% uranyl acetate solution increases the contrast. Panel A-D show central zoom outs from the region displayed in panel A. A large layer of listeriolysin O assembly is visible. Pores assembled on supported lipid bilayers have a comparable density (19).

### B. Listeriolysin O and Pneumolysin Assembly on a Lipid Monolayer

The physiological mechanism of listeriolysin O and pneumolysin is to assemble to a ring-shaped complex on a bilayer membrane. The ring inserts itself and forms a pore (18; 20). What would happen when the protein is prepared in the same way as it is done for an intra-membraneous protein, spraying it on-top of a lipid monolayer? Figure 8 shows the result.

Some ring-like structures of the expected size are visible, but the efficiency is dramatically reduced in comparison to the preparation on a lipid bi-layer. This is in agreement with the mechanism how listeriolysin and pneumolysin pores form. The experiment shows again that nanoelectrospray is indeed able to prepare lipid mono- and bi-layer structures which behave radically different when interacting with membrane proteins or protein complexes.

### C. Direct Observation of Protein Complex Formation

All observations reported so far were systematic results, repeated several times. Here, a sporadic observation is shown that supports the interpretation that the electron dense centers of the ring like structures are glycerol filled pores and that highlights the potential of the nanoelectrospray based preparation methods: the direct observation of the pore formation process of pneumolysine, from monomer to completed pore.

The images in figure 10 come from a preparation of pneumolysin on lipid monolayers. Initially hardly any pores should have formed (see figure 9). However in this case the spray was not operated at the desired low flow rate. Larger droplets were generated which landed on the surface membrane and the water evaporated leaving crystalline salt residuals on the surface. Nanoelectrospray ion sources on mass spectrometers operate with volatile buffers but for these experiments a more conservative approach was taken and the original solution the proteins were kept in was not changed. After the preparation was finished enough lipid was sprayed to form bi-layers on the surface. The images in figure 10 suggest that during incubation time pneumolysin monomers slowly dissolve from the crystalline residue and diffuse out into the glycerol layer. Because they encounter a lipid bi-layer they start to assemble and form pores. The entire process is visible and suggests that it proceeds sufficiently slow to be stretched out over the diffusion range. Glycerol has a significantly higher viscosity than water. And - diffusion in thin layers is slower than in bulk solution. All this suggests that protein complex formation might be directly observable in thin glycerol layers in particular when the temperature of the layer is controlled.

**Fig. 9.**
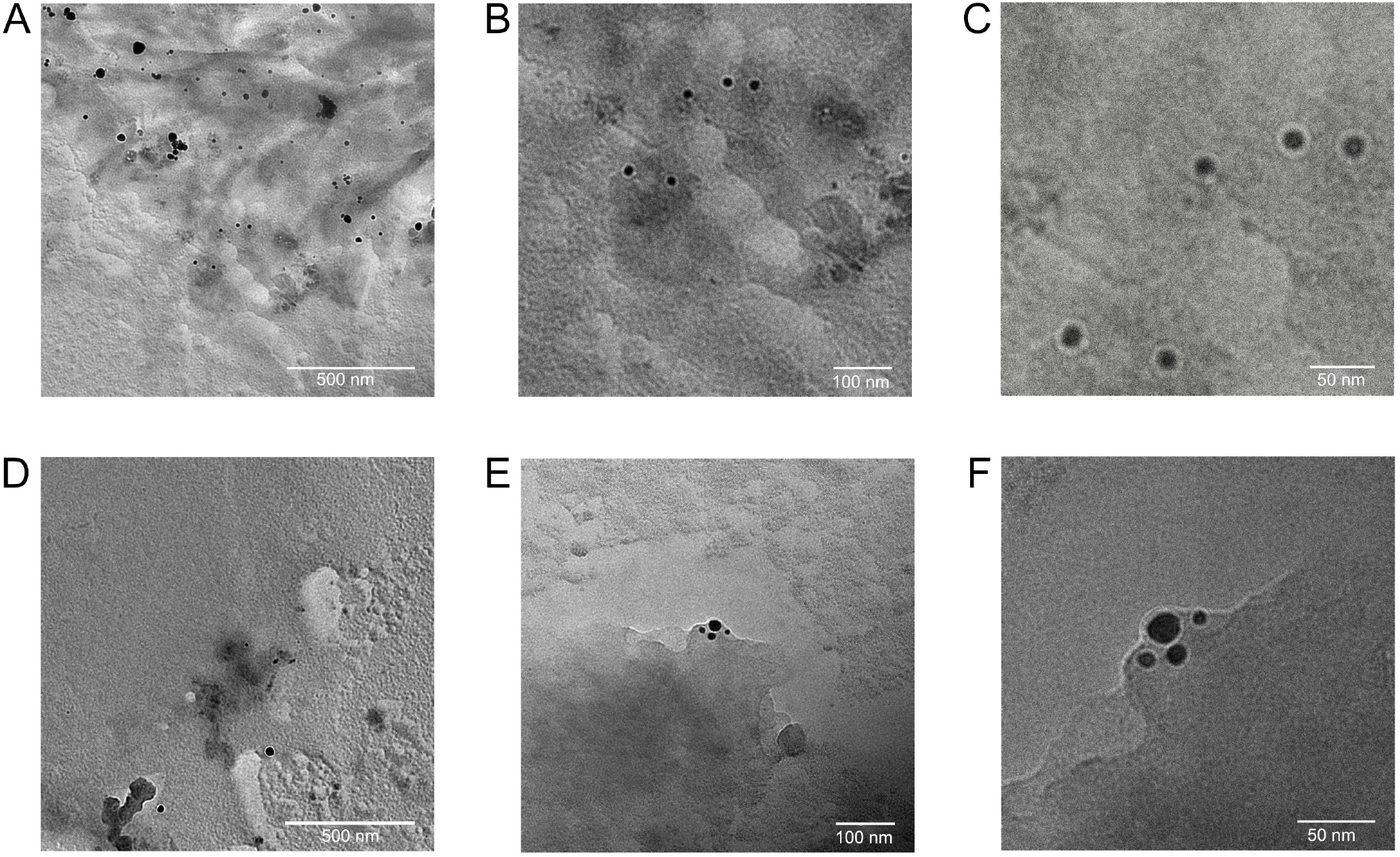
Listeriolysin O and Pneumolysin on a Lipid Monolayer. Pneumolysin and Listeriolysin O sprayed onto a lipid monolayer: panels A, B, and C show a pneumolysin preparation, panel D, E, and F a listeriolysin O experiment. The samples were shadowed with a 1.5 nm thick platinum layer at an angle of 15°.

**Fig. 10.**
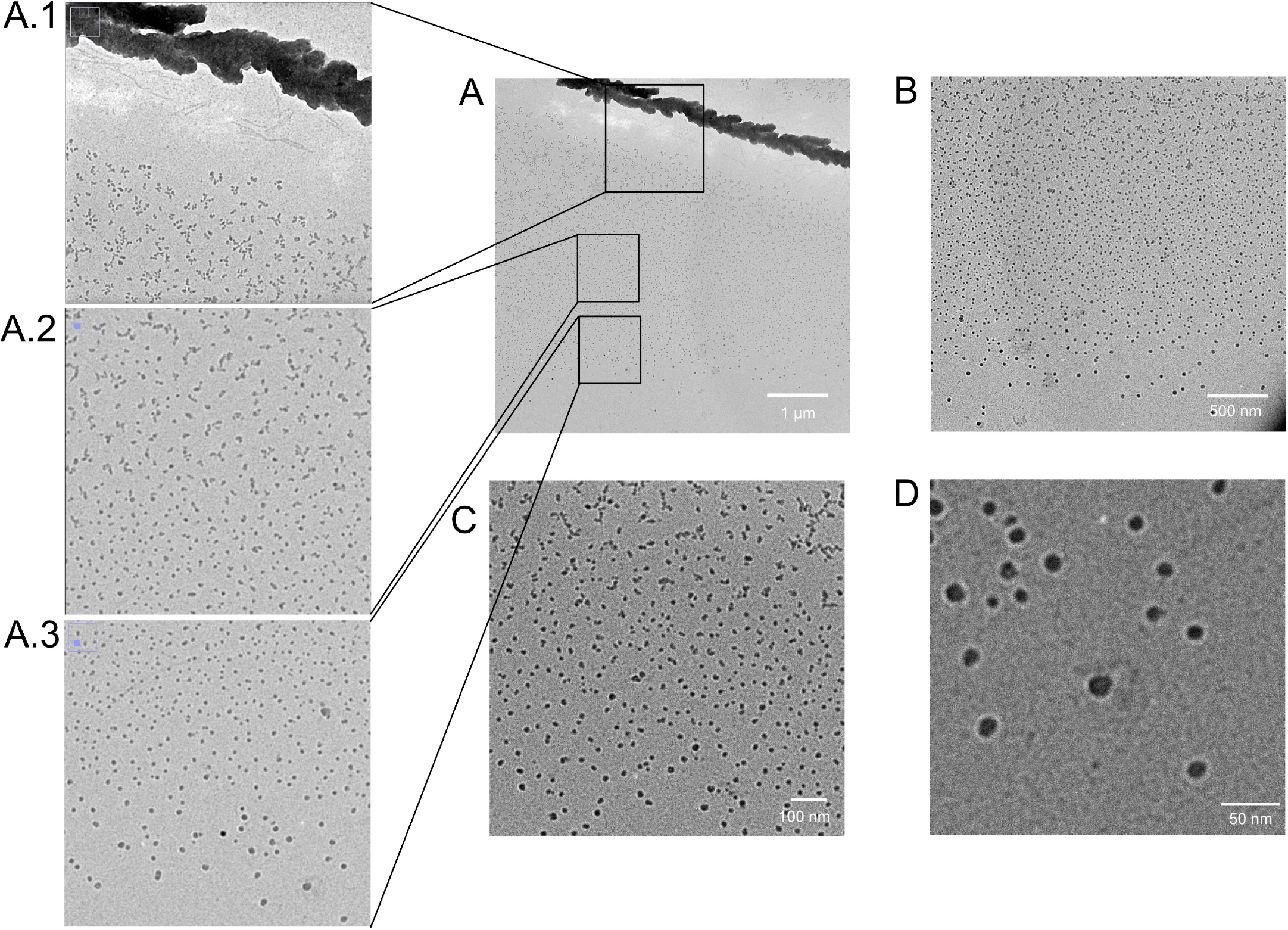
Direct Observation of Pneumolysin Complex Formation. Pneumolysin preparation on a lipid monolayer: pneumolysin was sprayed onto a lipid monolayer and covered with glycerol. After five days incubation, the layer was transferred to a carbon coated grid, washed with buffer, negatively stained with 1% uranyl-acetate and washed with water.

A pore formation in a supported bilayer has been directly observed using atomic force microscopy for the membrane attack complex (MAC) (21). The pore completes in about 100 seconds. This experiment here suggests that in glycerol layers it is even slower.

## V. Conclusions

The second of the two original hypotheses that nanoelectrospray should allow the composition of surfaces on a molecular level appears to hold. The fact that it was possible to self-assemble a large ompG containing membrane when a lipid monolayer acts as base but not with a lipid bilayer confirms this hypothesis. The same is true for the formation of listeriolysin O and pneumolysin pores. Only when sprayed upon a lipid bi-layer pore formation proceeds effectively.

These experiments describe a way how to generate large, protein containing membranes which are freely transportable. So far, the experiments did not aim at establishing an anisotropic orientation of the proteins. However, the proteins are likely to become oriented when the membrane assembly process proceeds in a static electric field.

It is very desirable to further undermine the conclusions that large membranes with integrated membrane proteins had been generated. Light microscopic images could help to prove that the membrane proteins are truly present and to capture the entire extend of the membrane. Functional experiments should support that the proteins are biologically active. And the technique should be tried with other membrane proteins to prove its genuine character. All of this is addressable when the electrospray technique is embedded into an appropriate environment. Even though all these experiments might be necessary to prove that membranes with embedded proteins had indeed been generated, this interpretation remains the most likely explanation for the observed experimental data.

## VI. Materials & Methods

### A. Protein Expression & Purification

Yildiz et al. describe the expression and purification of the outer membrane porin G (ompG) (11). Briefly, E. coli expresses the amino acids 22 - 301 plus an N-terminal methionine of the OmpG gene having cloning it into a pET26B plasmid vector. Unfolded ompG elutes from an ion exchange column after breaking the cells, centrifuging and solubilizing them in 8 M urea, 1% Triton-x 100 and 25 mM Tris-HCl. The refolding of the protein occurs in a 1% (weight/ vol) n-octyl-β-D-glucopyranoside (OG), 3 M urea solution. A final ion-exchange separation succeeds in removing unfolded or partially folded protein from the solution. With ultrafiltration or dialysis using membranes with a 12 kDa weight cutoff against a buffer containing 20% PEG-35000 the protein concentration rises to approx. 50 mg/ml. A buffer containing 10 mM Tris, 1% octylglucoside, 20 pmol/μl phosphatidylcholine with a concentration of 32 pmol/μl ompG serves as the base-solution to prepare ompG layers. The final spraying solution contains an additional 25% glycerol to protect the protein from the high charges of the electrospray droplets.

Listeriolysin O (LLO) came from two sources. LLO was expressed and purified as described in (16). Briefly, for expression in E.Coli BL21 (DE3) the gene segment encoding residues 25 - 529, the LLO sequence without the N-terminal secretion signal, complemented by an N-terminal His6 tag was cloned into the plasmid pET15b. The cells were grown overnight and then transferred into a selective medium containing 100 μg/ml ampicillin. After reaching a cellular density A600 of 1.2, the reduction of the temperature from 37°C to 30°C reduced the proliferation of the cells. The addition of 1 mM isopropyl β-D-1-thiogalactopyranoside inhibited the plasmid’s Lac-repressor and the protein expression starts. After 4 hours, the cells were collected, suspended in 50 mM Tris pH 7.7, 150 mM NaCl, and disrupted using a microfluidizer (M-110L, Microfluidics Corp., MA). After 1 hour centrifugation at 12 000 g, the supernatant was loaded onto a Ni-column to extract the His6-tagged protein. The protein was cleaved from the column with thrombin, concentrated on a 30 kDa cut-off membrane and purified with a gel filtration column (Superdex200) using 25 mM Tris (pH 7.7), 150 mM NaCl as running buffer. Recombinant LLO from Abcam (Cambridge, UK) was an alternative source for the protein (residues 60 - 529, product code ab83345).

The method to obtain pneumolysin (PLY) was similar (20). An Escherichia Coli colony expressed the His6 tagged version of the protein by cloning it into the plasmid pET15b. The cell culture grew overnight at 37°C. Transfer to a selective medium containing 50 μg/ml ampicillin ensured that only plasmid containing cells survived. With reaching an A600 between 1 and 1.5, the temperature is reduced to 30° to reduce cell proliferation. The addition of 0.5 mM isopropyl β-D-1-thiogalactopyranoside induced protein expression. After overnight expression, the cells were harvested, suspended in lysis buffer and disrupted using a microfluidizer. It followed the centrifugation of the cells for 1 hour at 185 000 g and loading of the supernatant onto a HisTrap FF column. With a lysis buffer containing 500 mM imidazole to outcompete the histidine-tag from its Ni+ binding sites, the protein elutes from the column. Using a 50 mM Tris pH 7.0, 5 mM β - mercaptoethanol solution, the protein fractions are diluted. Overnight incubation with 100 unit Thrombin at 4°C removed the His tag. Passing the protein solution over an HiTrap Q FF using a salt gradient from 0 to 1 M NaCl in a 50 mM Tris, pH 7.0, 5 mM β - mercaptoethanol buffer resulted in a purified sample. The protein solution was not frozen but kept at 4°C to avoid precipitation.

### B. Static Secondary Ion Mass Spectrometry

Electrospray prepared targets were investigated using Time-Of-Flight Secondary Ion Mass Spectrometry (TOF-SIMS) I and II instruments (9; 22; 23). The TOF-SIMS I instrument had a pulsed Ar+ ion source. The ions hit a grounded metal target. 8 keV Ar+ ions were focused onto a 1 mm spot with a primary ion density of 1010 Ar+ ions per (cm2·second) (static SIMS conditions). Secondary ions were extracted from the grounded target and left to drift into two time-of-flight regions connected by a 163° angular and energy focusing toroidal Poschenrieder condenser lens. A single ion counting device detected analytical ions. They were post-accelerated before being converted to electrons in a channel plate which excited a scintillator. A photomultiplier counted the photon-pulses. For the TOF SIMS II instrument, the Poschenrieder lens was replaced by an energy compensating reflectron which increased the mass resolution to more than 10 000. Significant for this investigation were improvements in the primary Ar+ ion source. The ion current was more stable than on the TOF-SIMS I instrument, and the diameter of the ion beam on the target was 0.1mm. The, for the time, small focus made spacially resolved scans of the prepared target regions possible.

### C. Electron Microscopy

Carbon-coated electron microscopic grids were briefly exposed to gas discharge to render them hydrophilic. Lowering a grid carefully onto a liquid meniscus carrying the prepared layers transfers them to the electron microscopy target. Several exchanges of double distilled water washed the specimens extensively. For inspection in the electron microscope, the grids were negatively stained using 1 % wt/ vol uranyl acetate. Alternatively, a platinum/carbon beam shot at an angle of 15° covered the surface with a metallic layer of up to 1.5 nm. An FEI Tecnai Spirit BioTWIN transmission electron microscope operating at 120 kV with varying magnifications of up to 150 000x was used for inspection.

### D. Electrospray based Surface Preparation

#### Electrospray Apparatus

An electrospray apparatus was built to distribute dissolved material evenly over a flat surface (7) (second generation instrument see figure 1). The first instrument had a metal nozzle with an emitter diameter of 250 μm. The balance between the electrostatic pull on the liquid and the flow resistance of the opening regulated the flow rate. The flow resistance was adjustable by varying the height of a precisely fitted metal cylinder within the outlet. Like this, flow rates between 8 and 15 μl/min were achievable. After the initial experiments, the nozzle was replaced by a hand-pulled, gold-coated glass capillary to reduce the stable flow rate to below 500 nanoliters/min (7). The liquid sample was sprayed from the reservoir under atmospheric conditions onto a flat metal target. By applying a high voltage, the tip of a liquid Taylor Cone, formed at the nozzle, emitted a stream of rapidly spreading, highly charged small droplets. The volatile solvent evaporates in flight, and the dried down residues cover the target region in a circle with a diameter of about 4 mm. The distance between the nozzle and the target is adjustable. For a second generation of the apparatus, the glass capillaries were manufactured by a commercial capillary puller (model P-87 Puller, Sutter Instruments Company, Novato, CA, USA) and mounted in a pressurized container (figure 1). Before spraying, the narrow glass tip was broken to have an orifice of about 1 μm in diameter (24). With this more mature device, the electrospray operates with stable and constant flow rates between 20 and 100 nanoliters/min.

#### Surface Preparation

For the investigation with the static Secondary Ion Time-of-Flight Mass Spectrometer (TOF-SIMS) a rhodamine/acetone solution was sprayed onto an aluminum target.

The target for the membrane preparation was the liquid meniscus in a small cylindrical container with a diameter of 3 mm filled with a 10 mM Tris solution, pH 7.5 which contained some SM-2 Biobeads (Bio-Rad, CA, USA). Egg-phosphatidylcholine was from Sigma-Aldrich. The final protocol for the membrane synthesis consists of five steps. The first material sprayed onto the buffer is the equivalent amount of a lipid bilayer of bipolar lipids dissolved in ethanol: 1 μl 66 pmol/μl lipid with 35% cholesterol, 65% phosphatidylcholine. This preparation is incubated overnight at room temperature. In a second step, a thin glycerol layer is added by spraying a 1:2 solution of glycerol/ethanol onto the meniscus for about 10 minutes. The third step consists of spraying enough lipid to generate half a bilayer: 1 μl 33 pmol/ μl lipid with 35% cholesterol, 65% phosphatidylcholine in ethanol. In the fourth step, the sprayed membrane protein and lipid complete the molecular mix to allow the self-assembly of an intact, the protein containing lipid bilayer. The sprayed solution consisted to 25% of glycerol to protect the protein from the high droplet charges when the water evaporates in flight: 1 μl ~32 pmol/μl ompG, 20 pmol/μl lipid 35% cholesterol, 65% phosphatidylcholine, 1% octylglucoside in 10 mM Tris, 25% glycerol. As the last step, another glycerol layer is added to complete a hydrophilic environment: 1:2 glycerol/ethanol solution for about 10 minutes. The entire assembly is incubated for 6 days at room temperature to allow for the extraction of the octylglucoside detergent by the SM-2 beads from the surface layers.

The buffer in the container for the pneumolysin and listeriolysin O preparation was a 50 mM Tris, 150 mM NaCl aqueous solution. For the listeriolysin O solution, the pH was adjusted to 5.5, for of the pneumolysin solution to 7.0. The first step in generating a protein membrane layer is to spray 1 μl of 66 pmol/μl lipid solution with 35% cholesterol and 65% phosphatidylcholine in ethanol to allow the formation of a closed lipid bilayer by overnight incubation. A thin glycerol layer generated by spraying a 1:2 solution of glycerol/ethanol for approximately 10 minutes creates a hydrophilic base for the subsequent membrane assembly.

The next step consists of adding either a half or a complete lipid bilayer. As before, incubation overnight completes bilayer formation. Now, the surface is prepared to add the protein containing film. The pneumolysin solution consisted of 26 pmol/μl protein in 50 mM Tris, 150 mM NaCl at neutral pH and the listeriolysin O solution had a concentration of 20 pmol/μl dissolved in 50 mM Tris, 150 mM NaCl at ph 5.5. The solution sprayed had an additional component of 25% glycerol. When the surface carried only a lipid monolayer, the protein solution contributes enough lipid to complete the bilayer - 20 pmol/μl lipid, 35% cholesterol, and 65% phosphatidylcholine. About 1 μl of the final solution distributed over the 3 mm large liquid meniscus generates the protein containing layer. Spraying a 1:2 glycerol/ethanol solution for 10 minutes finalizes the hydrophilic environment. The self-assembly process proceeds throughout the incubation period of several days at room temperature.

## Supporting information

Overlay simulation of ompG crystal structure

## Authors Contribution

M.W. did all the experiments and calculations mentioned in the article. The proteins and lipids were contributed by the groups of Prof. Kühlbrandt and Dr. Oezkan Yildiz.

## Acknowledgements

All experiments were done in the laboratory of Prof. W. Kühlbrandt at the Max Planck Institute of Biophysics, Frankfurt, Germany. I am grateful to Prof. Kühlbrandt for providing the necessary support to conduct all these experiments. I thank Dr. Oezkan Yildiz and Dr. Katharina von Pee for the intellectual and material support. Without the help of Deryck Mills on the electron microscope and with technical questions, the project could never have been done.

The work of Matthias Wilm was supported by the Science Foundation Ireland, SFI grant 07/SK/B1184c.

## VII. Supplemental Material

**Fig. I.**
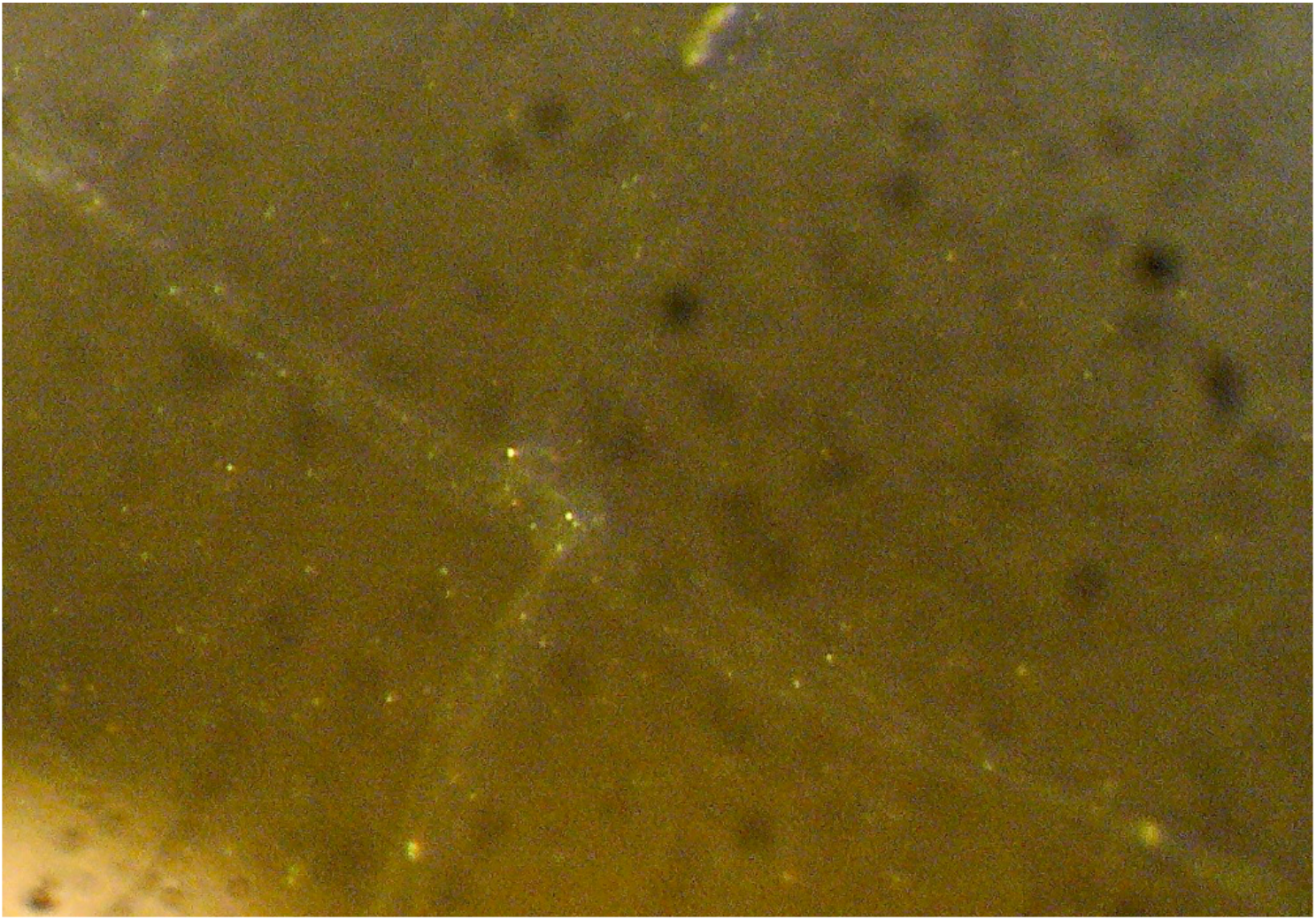
Lipid Bi-Layer on Buffer. Stereomicroscopic image of a nanoelectrospray generated lipid bilayer on a buffer solution after overnight incubation at room temperature. Some folds of the viscous surface are visible.

> **<u>OmpG-Simulation.mp4</u>**

**Video I Overlay of the Crystal Structure of ompG and an ompG Containing Membrane**

Overlay of an ompG crystal structure derived image (from Behlau, M., D. J. Mills, H. Quader, W. Kühlbrandt & J. Vonck. 2001. Projection structure of the monomeric porin OmpG at 6 A resolution. *Journal of molecular biology* 305:71-77) with the platinum shadowed transmission electron image of the membrane containing ompG (figure 7). The scales of both images have been matched.

